# Anastasis induced by bee venom in normal cells compared to persistent cell death in breast cancer cells

**DOI:** 10.1101/2024.02.18.579605

**Authors:** Sinan Tetikoğlu, Ugur Uzuner, Selcen Çelik Uzuner

## Abstract

Anastasis is a phenomenon that has been recently defined as a return from induced apoptosis. Its mechanism has not been clearly elucidated. Anastasis is thought to be involved in the development of drug resistance in cancer cells, however the distinct regulation of anastasis in normal and cancerous cells during anti-cancer therapy has not been discovered. One of the most privileged therapy strategies focuses on the drugs that are selectively cytotoxic in cancer cells but not negatively affect normal cell proliferation. This study for the first time comparatively evaluated the anastatic effect of a common synthetic cytotoxic agent, cisplatin and a natural cytotoxic agent, bee venom. The study showed that bee venom induced anastasis in normal cells (MCF10A, NIH3T3 and ARPE19) but cancer cells (MDA-MB-231 and MCF7) were irreversibly in cell death process. Liver cancer cells (HEPG2) were more resistant to bee venom-induced persistent cell death and tended to recover at higher concentrations compared to breast cancer cells. However, cisplatin induced persistent cell death in both normal and cancerous cells. Besides, selectivity indexes of bee venom in terms of IC50 values were higher than cisplatin. This study indicates that bee venom produces such an effect by selectively inducing anastasis only in normal cells suggesting that bee venom has prominent potential for cancer therapy, in particular for breast cancer, with recovery and maintenance of normal cells’ viability.

**Graphical Abstract:** 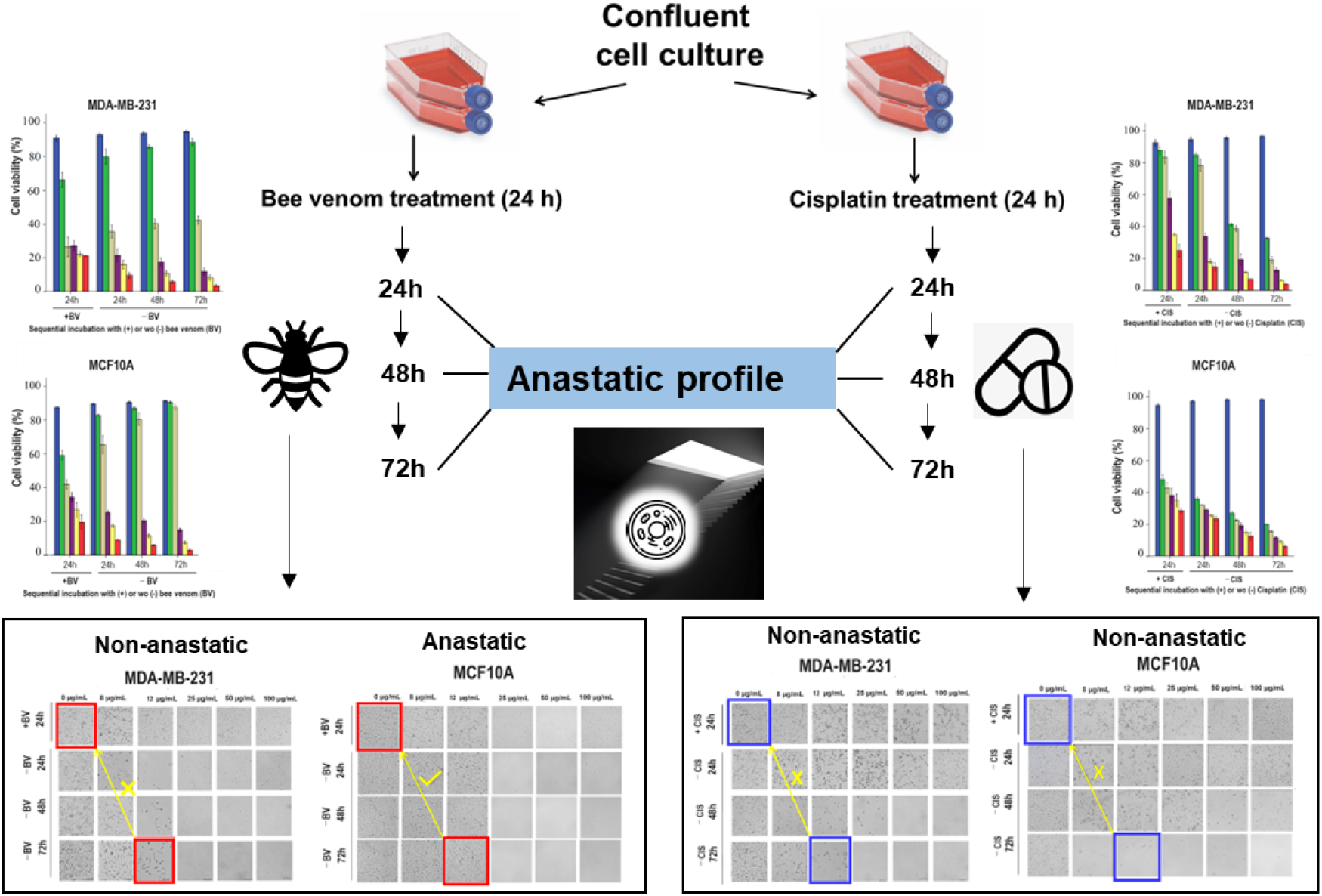

## Introduction

Anastasis is a mechanism defined as a reversal of apoptosis in the cells followed by recovery. This phenomenon, first described in 2012, showed that cell death processes are reversible after treatment of different cell types with ethanol, and this reversal was named as ‘anastasis’. After the cells were incubated with ethanol for a certain period of time followed by the removal of ethanol, anastasis was observed in the cells (H. M. Tang et al., 2017, 2022; H. M. Tang & Tang, 2018; Ho Lam Tang et al., 2012). There is evidence that the anastasis mechanism is different from another cell survival mechanism such as autophagy (Sun et al., 2017). Additionally, some genes predicted to be involved in the anastasis process have been reported (H. M. Tang et al., 2022). However, the cellular pathways for anastasis mechanism have not been fully elucidated.

Anastasis appears to be an effective mechanism for triggering survival of apoptotic cells. Recently, the development of resistance of cancer cells in chemotherapy treatment processes suggests that cancer cells can escape apoptosis by activating anastasis (Mohammed et al., 2022). From this perspective, the anastasis event is of great importance in terms of explaining the developmental mechanisms that play a role in cancer recurrence. However, the potentials of anti-cancer agents for anastasis in normal cells while apoptosis in cancer cells remain unrevealed. Therapeutic agents for cancer treatments are expected to have a high cytotoxic effect on cancer cells while showing less cytotoxicity in healthy cells. Bee venom is a promising natural mixture to induce cytotoxicity in cancer cells (Abdulmalek et al., 2022; Jeong et al., 2019; Lim et al., 2019). Its cytotoxic function is prominenced as bee venom can selectively kill cancer cells (Sharkawi et al., 2015; Tu et al., 2008). Besides, a wide range of therapeutic potentials such anti-rheumatoid, anti-inflammatory and anti-aging activities of bee venom have been reported (El-Tedawy et al., 2020; Sung et al., 2023). Although bee venom is well-known to have a strong cytotoxic potential, its cytotoxic specificity regarding the discrimination between anastatic and non-anastatic profiles in cancer and normal cells has not been revealed. This study showed that bee venom significantly induced anastasis in normal cells while triggered irreversible cell death in breast cancer cells.

## Methods

### Cell culture

Cancer cells experienced in this study were 1) MDA-MB-231 (American Type Cell Collection ATCC, Cat No. HTB-26, VA, USA) human metastatic breast, 2) HEPG2 (ATCC, Cat. No. HB-8065) human liver and 3) MCF-7 (ATCC, Cat No. HTB-22) human breast cancer cells. Normal cells used as control were 1) ARPE-19 (ATCC, Catalog No. CRL-2302) human epithelial from retina, 2) MCF10A (ATCC, Catalog No. CRL-10318) human breast, and 3) NIH3T3 (ATCC, Cat. No. CRL-1658) embryonic mouse fibroblast. MCF-7/MDA-MB-231/ARPE-19, HEPG2 and NIH3T3/MCF10A cells were cultured in RPMI (Wisent Inc. Multicell, USA, Cat. No. 350000CL), EMEM (Cat. No. 320026CL) and DMEM (Cat. No. 319005CL) media, respectively. These media contained 10% (v/v) fetal bovine serum (Sigma-Aldrich Co. St. Louis, US, Cat. No: 12103C) for MCF-7/MDA-MB-231/ARPE-19/HEPG2/MCF10A cells and 10% (v/v) bovine calf serum (Sigma-Aldrich, Cat. No. 12133C) for NIH3T3 cells, and 1% (v/v) penicillin-streptomycin (Capricorn, Cat. No. PS-B) was added to each media. Cells were incubated at 37 °C incubator with 5% CO_2_. The passage numbers of these cells were used throughout the study were 19 for MCF-7, 8 for MDA-MB-231, 8 for HEPG2, 18 for ARPE19, 13 for MCF-10A, and 12 for NIH3T3.

### MTT cytotoxicity assay

5000 cells per well were cultured until they reached at full confluency followed by treatment with bee venom at 8, 12, 25, 50 or 100 μg/mL for 24h. Bee venom was obtained and prepared as previously (Çelik Uzuner et al., 2021). After 24h incubation with bee venom, cells were treated with 200μl media including 10 μL of MTT (3-[4,5-dimethylthiazol-2-yl]-2,5 diphenyltetrazolium bromide, 5mg/mL) (Sigma-Aldrich, U.S., Cat. No. M2272) per well for 4h at 37°C (Ayazoglu Demir et al., 2021; Çelik Uzuner et al., 2021). After incubation with MTT, 200 μL DMSO was added to each well and incubated for 2h on a shaker (in dark) at room temperature. After incubation, color development (from purple to yellow) was observed and absorbances were measured at 570 nm using a microplate reader. IC_50_ values for each cell line were calculated as previously according to logarithmic slope formula (Ayazoglu Demir et al., 2021). Each treatment was performed as at least three repeats and also each experiment was repeated independently as triplicates.

### Anastasis experimental design

All cells were treated separately with bee venom or cisplatin for 24h as stated above, and then washed once with 1x PBS (phosphate buffered saline) (Wisent, 311-010-CL). Cells were trypsinized and collected by centrifugation at 230 rpm. The supernatant was removed, and the cells were resuspended in fresh medium. Cell viability (%) and total live cell numbers were determined by trypan blue staining using Countess FL II automatic cell counter (Thermofisher) (Koç et al., 2018). 10μL of cell suspension was mixed (1:1 ratio) with 10μL of 0.4% trypan blue (Biological Industries, B 103-102-1B) and incubated for approximately 10 mins at room temperature. 10 μL of cell-dye mixture was loaded on the coverslip of the device. Trypan blue is a negatively charged dye that is used to determine cell viability. Since the membrane structure is intact in living cells, the dye cannot enter the cell, while dead cells absorb the dye and dead cells appear blue/black under the light microscope, and live/dead cells stained by trypan blue can be also detectable using automated counters. The rest of the cells were washed three times with 1x PBS to remove bee venom/cisplatin completely, and the washed cells were cultured again in fresh media without bee venom/cisplatin for 24 hours. The same method was repeated for each of the next 24 and 48 hours. The viability rates of the treated cells with bee venom/cisplatin in the first 24 hours were determined at the end of the next 24, 48 and 72 hours in the media that did not contain bee venom/cisplatin (**Figure 1**). It should be noted that all untreated cells proliferated too much therefore half of the cell suspensions were discarded after counting and seeded in fresh media.

**Figure 1.**
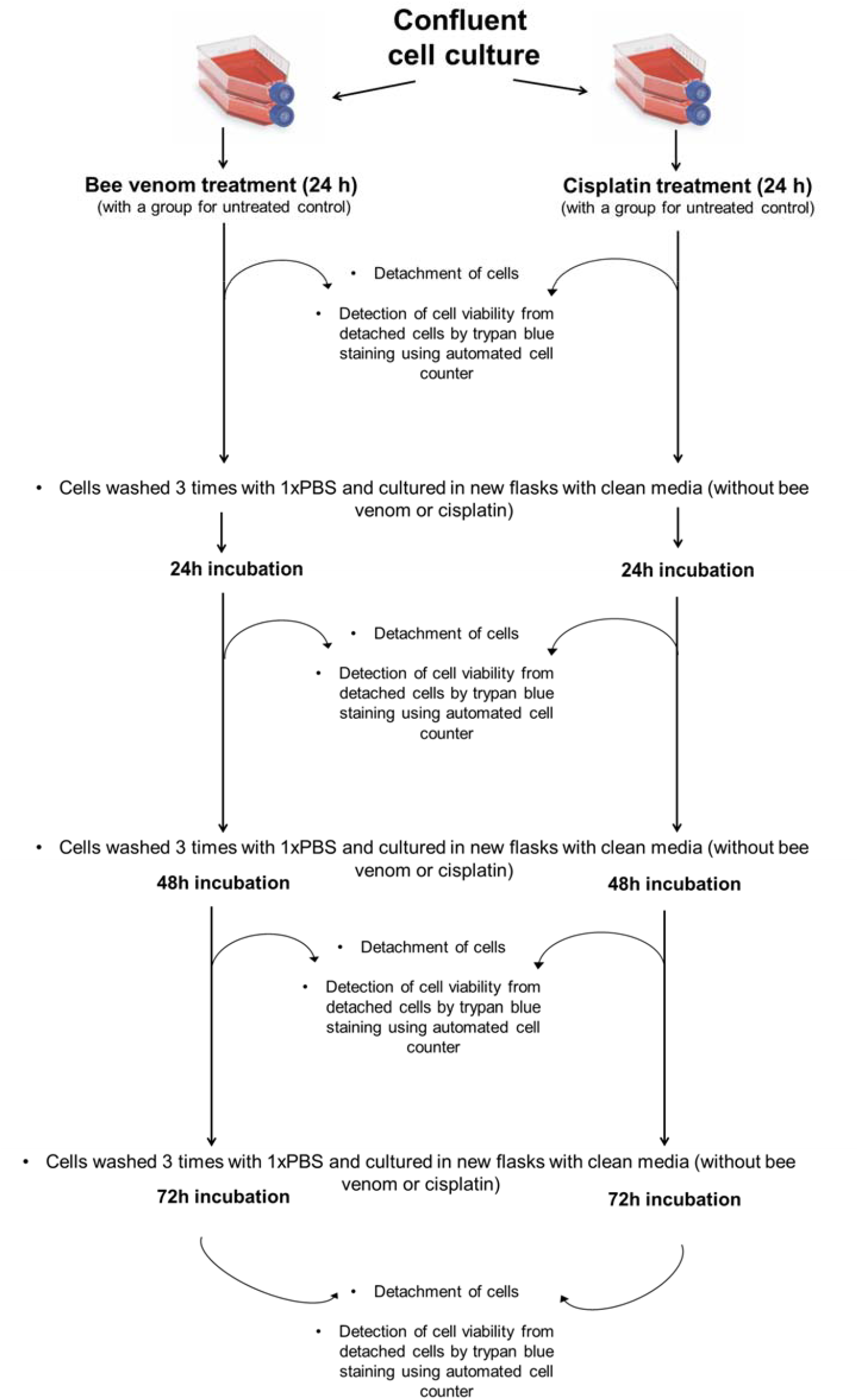
Experimental plan for anastasis.

### Population doubling times

Population doubling times were calculated using the formula below where *Nt* is the number of cells at time t, *N0* is the number of cells initially at time *0*, t is time (hours), and gr is the growth rate (AAT Bioquest, 2020).

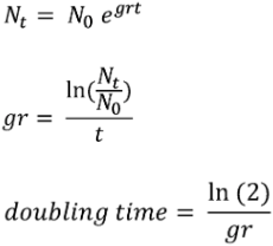

### Statistical analyses

All experiments were performed in at least three independent repeats. Each repeat also included at least three intra-experimental repeats for a treatment. The reliability of the automated counter was tested by measuring each sample at least three times within an experimental repeat. Standard errors of viabilities (S.E. +/-standard error of the mean) were calculated using SPSS software, and viabilities (%) were compared using UNIANOVA, and p values less than 0.05 were considered as significant.

## Results

First of all, to reveal cytotoxic doses for both bee venom and cisplatin, IC50 values were determined by MTT assay for each cell line after bee venom or cisplatin. Cancer cells were more sensitive to bee venom with IC50 values as 8μg/ml, 8μg/ml and 12μg/ml in MDA-MB-231, MCF7 and HEPG2 cells respectively (**Figure 2, left panel A-C**), however normal cells were more resistant to higher doses of bee venom with IC50 values as 36μg/ml, 38μg/ml and 50μg/ml in MCF10A, ARPE-19 and NIH3T3 cells, respectively (**Figure 2, left panel D-F**). Normal cells (except ARPE-19 cells) had a more tendency towards cell death after treatments with low doses of cisplatin, compared to bee venom. IC50 values of MDA-MB-231, MCF7 and HEPG2 cells were 12μg/ml, 20μg/ml and 12μg/ml, respectively (**Figure 2, right panel A-C**), and of MCF10A, ARPE-19 and NIH3T3 cells were 25μg/ml, 70μg/ml and 8μg/ml, respectively (**Figure 2, right panel D-F**). Changes in cell morphologies and population density of culture were shown by representative microscopy images after 24h incubation of bee venom or cisplatin at different doses (**Figure 2G**). In general, bee venom treatment resulted in higher selective indices compared to SI values detected after cisplatin treatment (**Table 1**). The higher SI suggests the higher selectivity for triggering apoptosis in cancer cells with a minimal cytotoxic effect on normal cells.

**Table 1.**
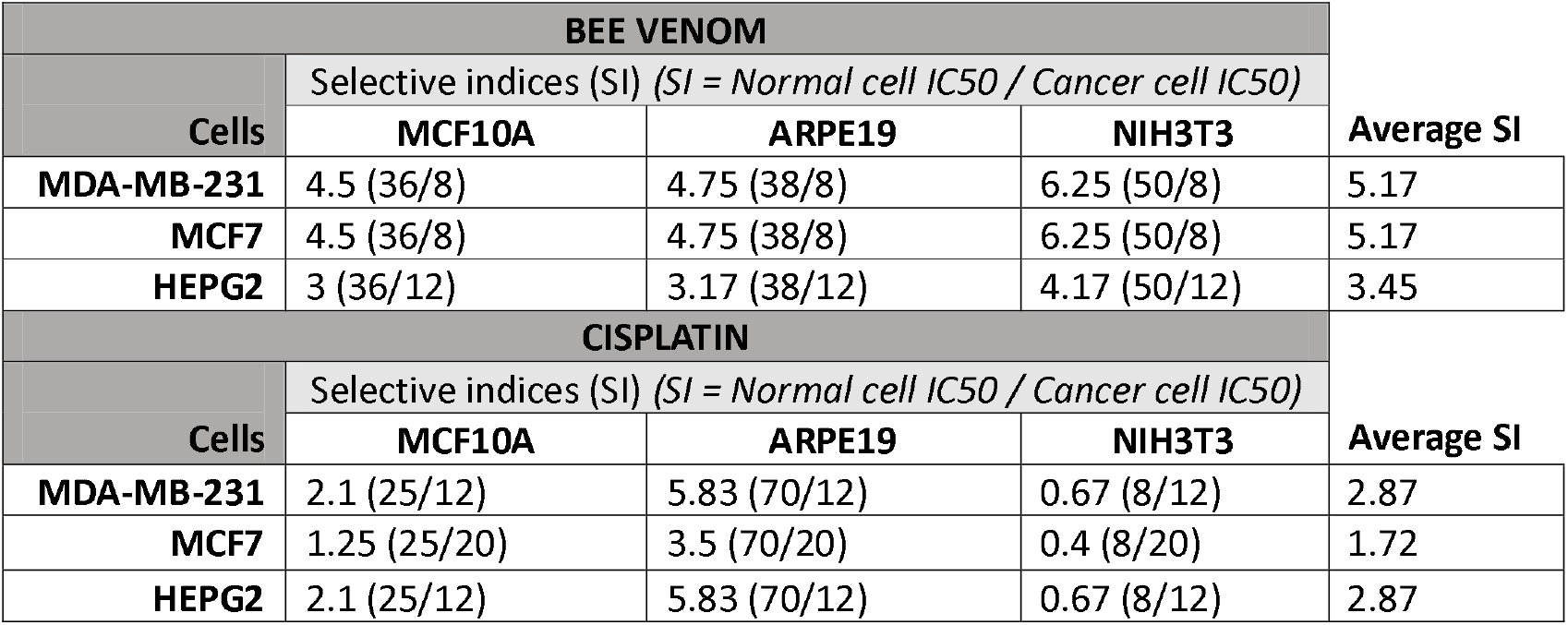
Selective index values (SI) after bee venom or cisplatin treatment.

| **BEE VENOM** |  |  |  |  |
| --- | --- | --- | --- | --- |
| **Cells** | Selective indices (SI) (*SI = Normal cell IC<sub>50</sub> / Cancer cell IC<sub>50</sub>*) |  |  | **Average SI** |
|  | **MCF10A** | **ARPE19** | **NIH3T3** |  |
| **MDA-MB-231** | 4.5 (36/8) | 4.75 (38/8) | 6.25 (50/8) | 5.17 |
| **MCF7** | 4.5 (36/8) | 4.75 (38/8) | 6.25 (50/8) | 5.17 |
| **HEPG2** | 3 (36/12) | 3.17 (38/12) | 4.17 (50/12) | 3.45 |
| **CISPLATIN** |  |  |  |  |
| **Cells** | Selective indices (SI) (*SI = Normal cell IC<sub>50</sub> / Cancer cell IC<sub>50</sub>*) |  |  | **Average SI** |
|  | **MCF10A** | **ARPE19** | **NIH3T3** |  |
| **MDA-MB-231** | 2.1 (25/12) | 5.83 (70/12) | 0.67 (8/12) | 2.87 |
| **MCF7** | 1.25 (25/20) | 3.5 (70/20) | 0.4 (8/20) | 1.72 |
| **HEPG2** | 2.1 (25/12) | 5.83 (70/12) | 0.67 (8/12) | 2.87 |

**Figure 2.**
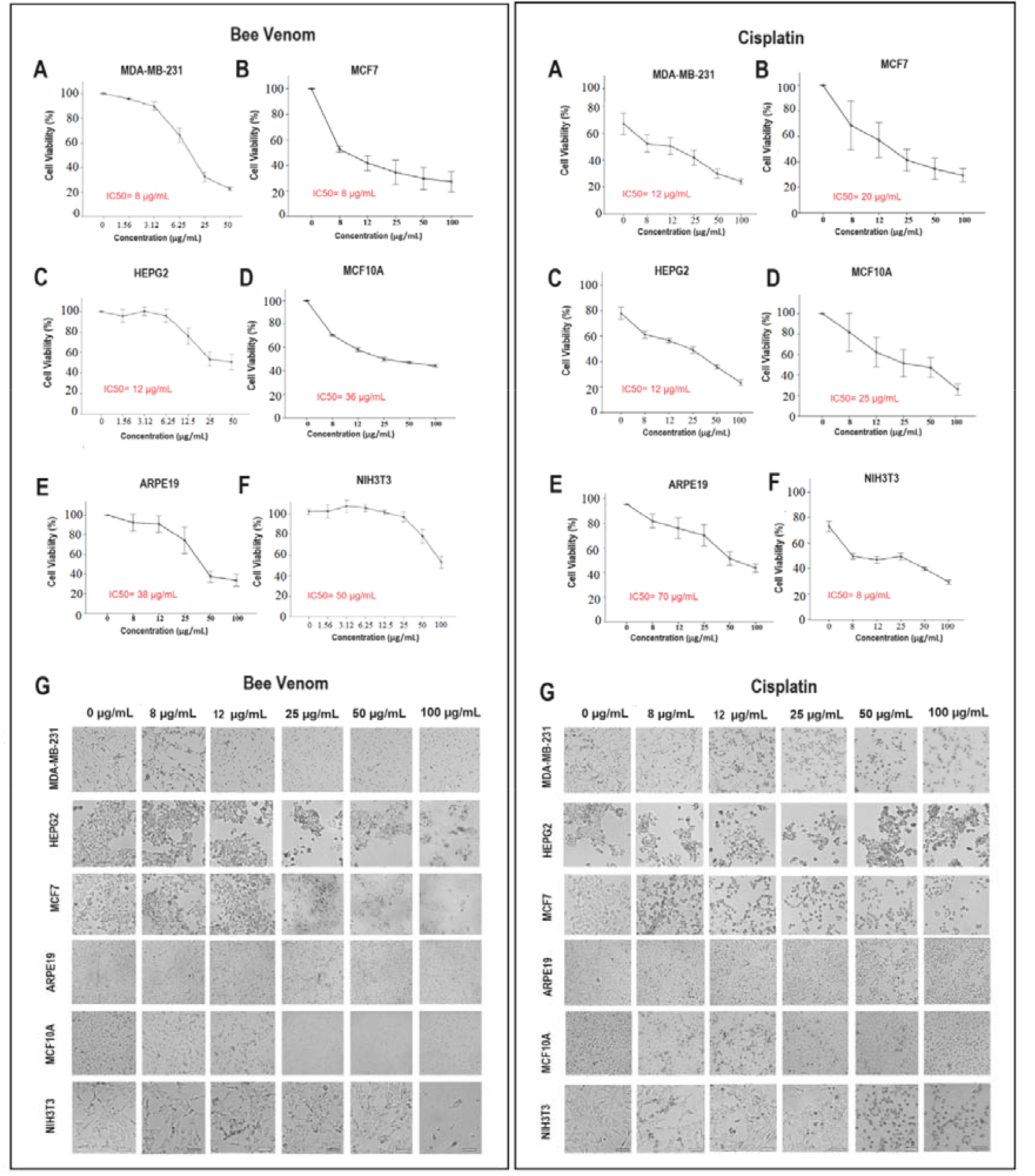
Cytotoxicity profiles in MDA-MB-231 (A), MCF7 (B), HEPG2 (C), MCF10A (D), ARPE19 (E) and NIH3T3 (F) cell lines after bee venom (left panel) and cisplatin (right panel)

We then assessed the anastatic response of cells against bee venom and cisplatin. To determine the numbers of living cells, MCF-7, MDA-MB-231, ARPE-19, HEPG2, NIH3T3 and MCF10A cells were treated with bee venom or cisplatin. A group of each cell was used as a control without any treatment. The cells were treated with agents, then washed and left for recovery up to 72h as mentioned in the method section (**Figure 1**). MDA-MB-231 cells continued to die even if bee venom (25 μg/ml and more) was removed, and they entered anti-anastasis after 12 μg/ml bee venom (**Figure 3A**). Another breast cancer cell line, MCF7, was sensitive to bee venom at each concentration after removal and each condition was anti-anastatic for MCF7 (**Figure 3B**). But HEPG2 tended to enter anastasis up to 25 μg/ml, and the minimum anti-anastatic dose of bee venom was 50 μg/ml in liver cancer cells (HEPG2) (**Figure 3C**). MCF10A cells recovered up to 12 μg/ml (**Figure 3D**), NIH3T3 cells were anastatic at each concentration including 100 μg/ml (**Figure 3E**), and ARPE-19 cells showed anastatic profile up to 25 μg/ml (Figure 3F). Statistical comparisons indicated that 12 μg/ml of bee venom was anastatic in normal cells (**Figure 4D-F**), however anti-anastatic in both breast cancer cells (MDA-MB-231 and MCF7) (Figure 4A, C). HEPG2 cells are more resistant to anti-anastasis since they have entered permanent cell death after 50 μg/ml (**Figure 3 and 4B**). The results suggest that bee venom significantly induced selective anastatic difference in normal breast cancer cells compared to two breast cancer cell lines. But none of the cells entered in the anastasis process after cisplatin (**Figure 5A-F and 6A-F**). Cisplatin was found to be selectively cytotoxic on HEPG2 cells (compared to non-cancerous epithelial cells, ARPE-19) with 5.83 SI value (**Figure 2, Table 1**), however cisplatin did not change anastatic response of ARPE19 cells compared to HEPG2 cells. This suggests that cisplatin also harms normal cells in long-term and normal cells cannot recover via induced anastasis.

**Figure 3.**
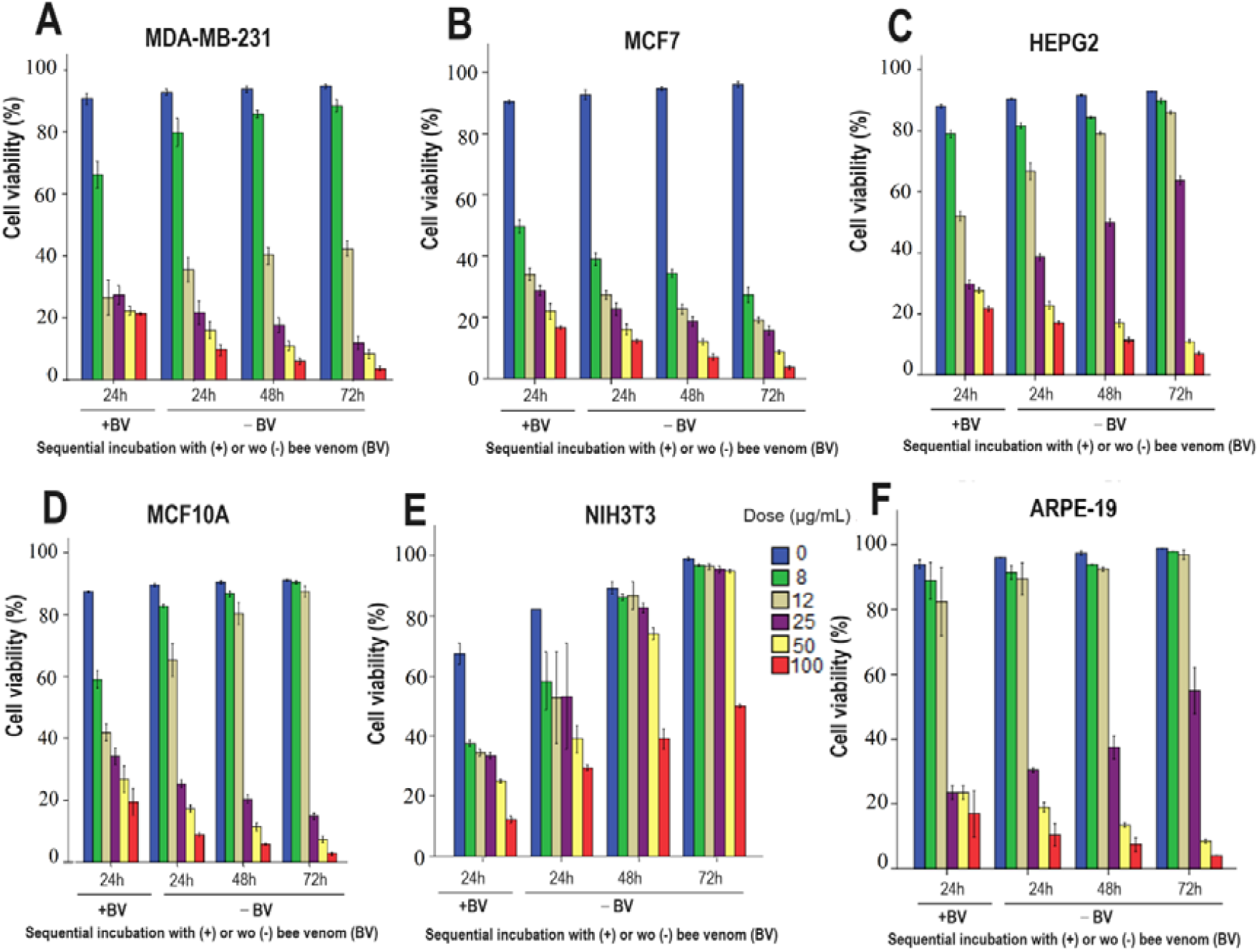
Bee-venom induced cell viabilities (%) in cancer (A, B, C) and normal (D, E, F) cells during anastasis.

**Figure 4.**
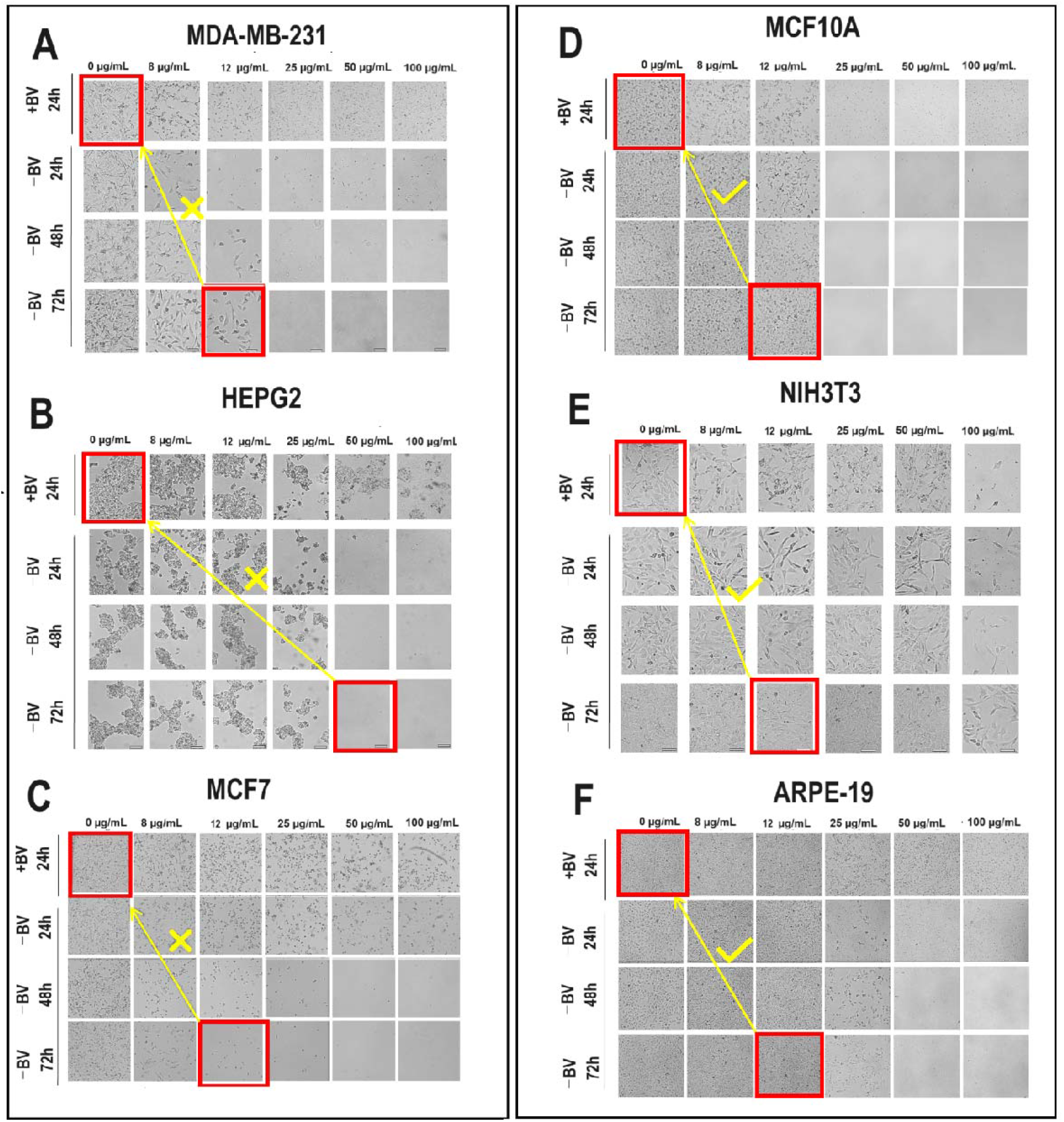
Bee venom-induced anti-anastatic profile in cancer (A, B, C) and anastatic profile in normal (D, E, F) cells.

**Figure 5.**
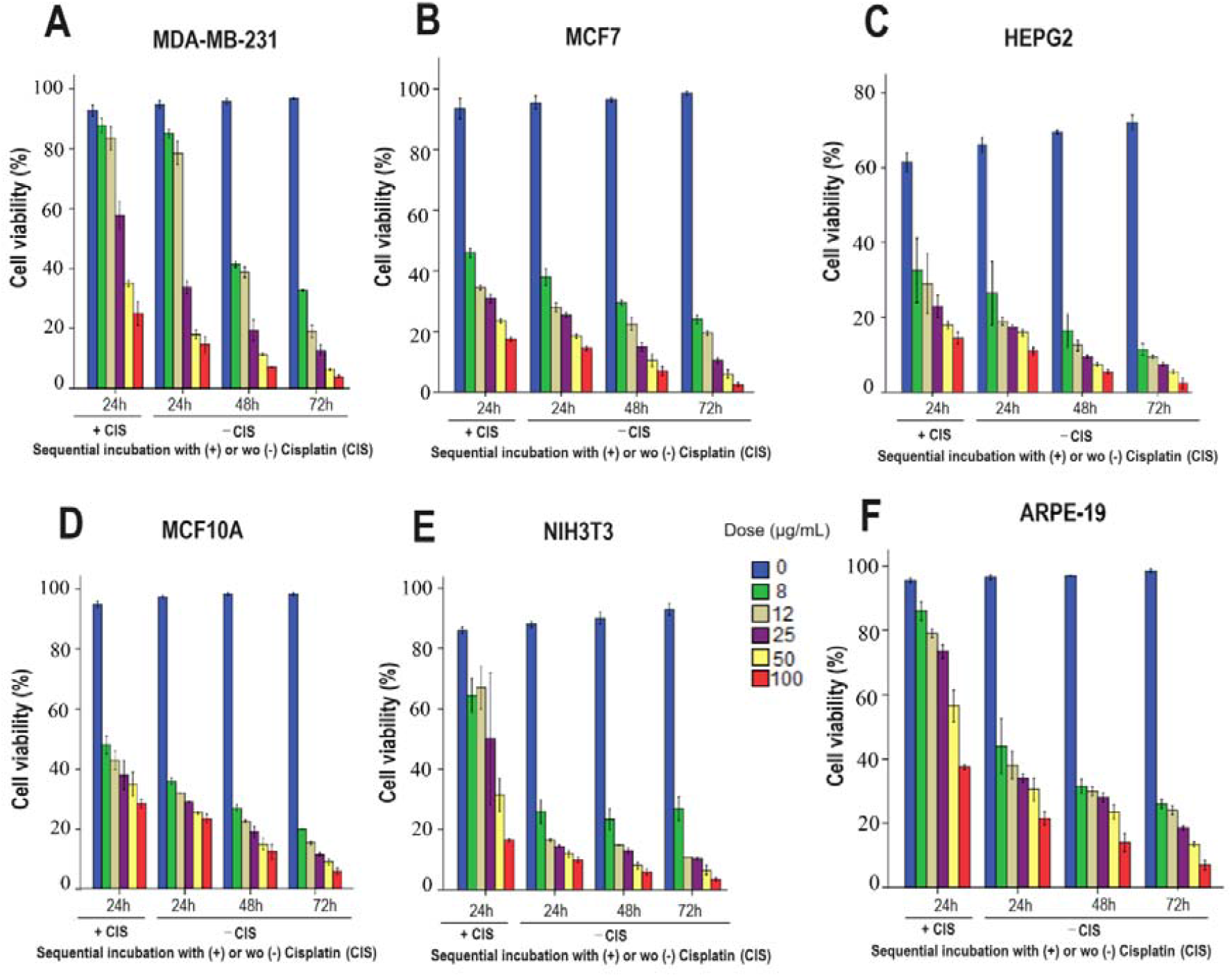
Cisplatin-induced cell viabilities (%) in cancer (A, B, C) and normal (D, E, F) cells in terms of anastatic expectation.

**Figure 6.**
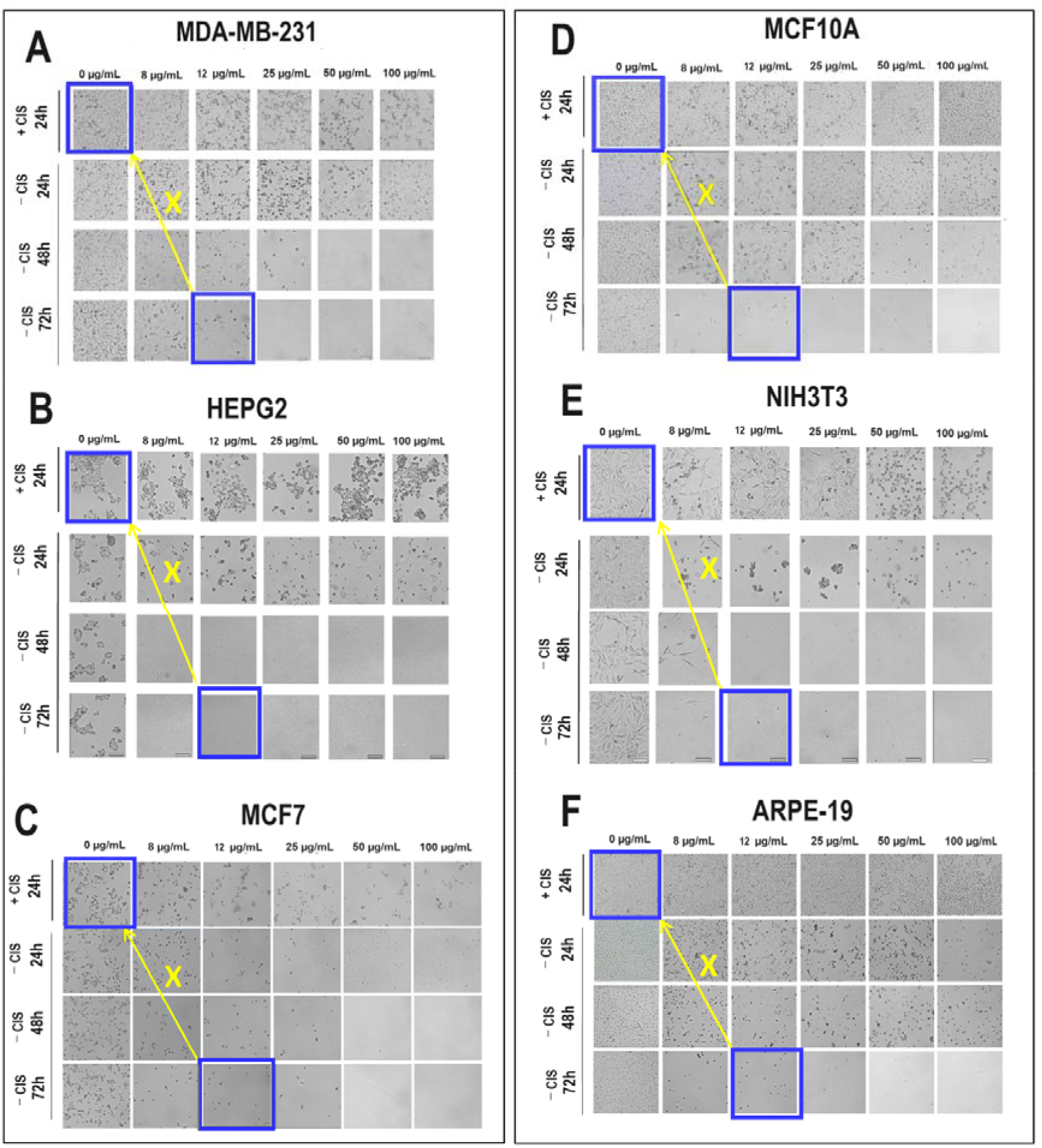
Cisplain-induced anti-anastatic profiles in cancer (A, B, C) and normal (D, E, F) cells, respectively.

Significant dose intervals that induced anastasis are summarised in **Table 2** for each cell line-agent combination. MCF7 was the most sensitive cancer cell line to both agents with no anastatic trend at any conditions. MDA-MB-231 cell line appeared to induce moderate anastasis only at the experienced lowest dose of bee venom (8μg/ml). Normal breast cancer cell line (MCF10A) was more anastatic to bee venom at higher doses compared to two breast cancers. However, epithelial liver cancer (HEPG2) was the most resistant cancer cell line to bee venom as these cells were anastatic up to 25 μg/ml which was similarly observed in normal epithelial ARPE-19 cells. The highest anastasis rate was observed in NIH3T3 cell line up to 100μg/ml.

**Table 2.** Significant anastatic dose interval in each cell line after bee venom or cisplatin (at 72h)

| Agent | Cancer cells |  |  | Non-cancerous cells |  |  |
| --- | --- | --- | --- | --- | --- | --- |
|  | MDA-MB-231 | MCF7 | HEPG2 | MCF10A | ARPE19 | NIH3T3 |
| Bee venom | 8 µg/ml | N/D | 8-25 µg/ml | 8-12 µg/ml | 8-25 µg/ml | 8-100 µg/ml |
| Cisplatin | N/D | N/D | N/D | N/D | N/D | N/D |

N/D, not detectable

We next analyzed population doubling times for each cell after different doses/incubations with bee venom or cisplatin. Live cell numbers were obtained during trypan blue staining and population doubling times were calculated using a formula given in the methods section. **Figure 7** shows each dose and incubation time in the cancerous (**A-C**) and in normal cells (**D-F**). **Table 3** indicates the average population doubling times between first incubation with agent (24h) and anastatic incubation for extra 72h.

**Table 3.** Average population doubling times (hour) (from the time for 24h incubation with bee venom to anastatic incubation for 72h) (N/D; not detectable)

| BV dose (µg/ml) | Cell Line |  |  |  |  |  |
| --- | --- | --- | --- | --- | --- | --- |
|  | MDA-MB-231 | HEPG2 | MCF7 | MCF10A | ARPE19 | NIH3T3 |
| 0 | 40,6676 | 37,5204 | 45,9007 | 65,2756 | 83,2831 | 38,6427 |
| **8** | 71,3494 | 63,1007 | N/D | 40,4933 | 68,4168 | 63,2968 |
| **12** | N/D | 51,4698 | N/D | 30,2763 | 74,8264 | 39,1825 |
| **25** | N/D | 162,155 | N/D | N/D | 40,6186 | 37,5684 |
| **50** | N/D | N/D | N/D | N/D | N/D | 38,4714 |
| **100** | N/D | N/D | N/D | N/D | N/D | 70,1518 |

**Figure 7.**
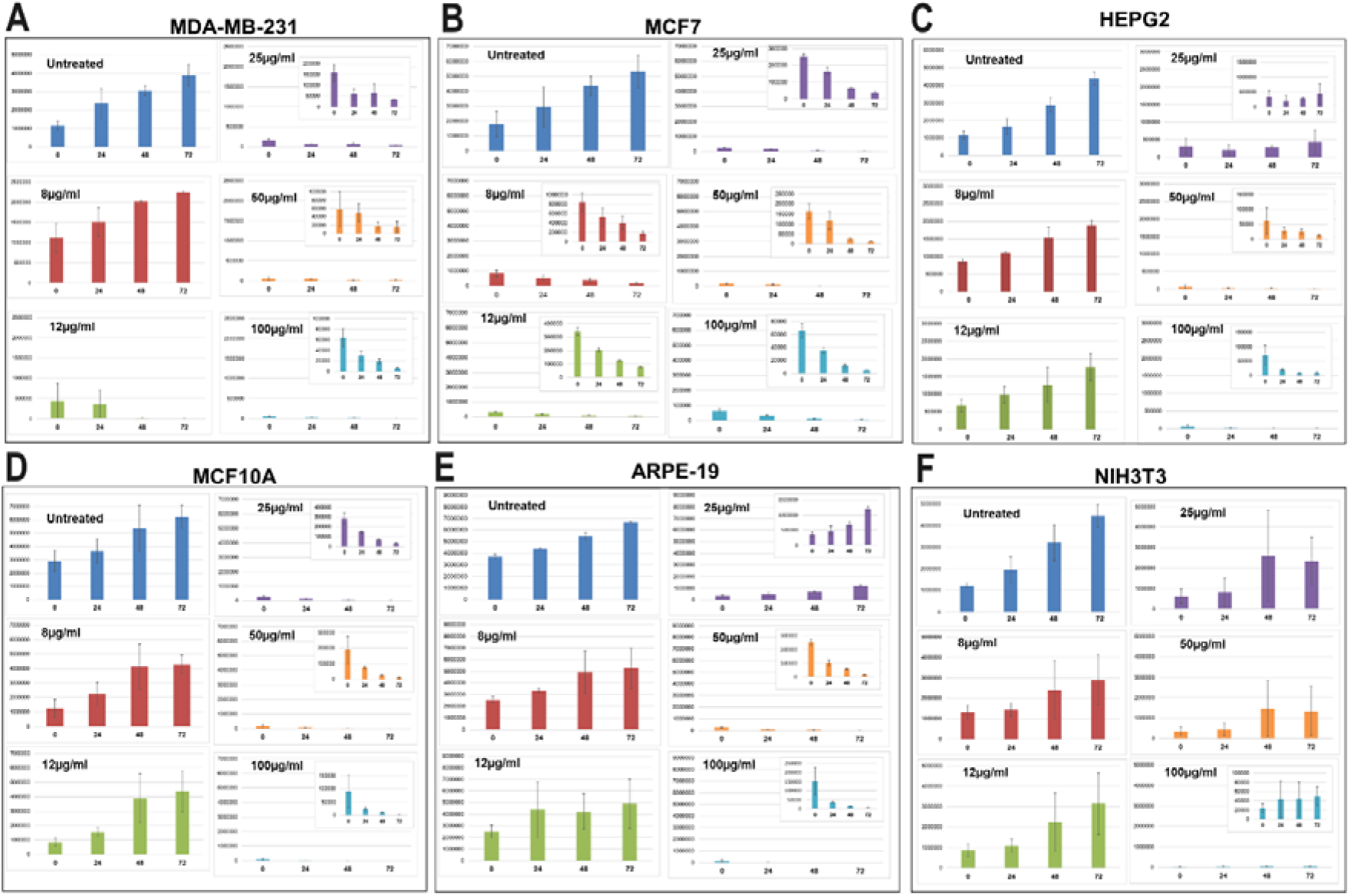
Average live cell numbers in cancer cells (A-C), and in normal cells (D-F) during anastasis-related response against bee venom.

Total live cell number increased at 8 μg/ml in MDA-MB-231 (**Figure 7A**) and 8-12 μg/ml in HEPG2 (**Figure 7C**), whereas live cell number did not increase at any doses in MCF7 cells (**Figure 7B**). HEPG2 cells are the only cancerous cells that have a tendency towards a degree of cell proliferation at 25 μg/ml. Most of MCF10A were progressively live in a time-dependent manner up to 12 μg/ml (**Figure 7D**), but both ARPE-19 and NIH3T3 cells were further alive at 25 μg/ml (**Figure 7E, F**). Population doubling times were consistent with live cell trend (**Table 3**). MCF7 was the most sensitive cancer cell line that could not recover via anastasis with a constant decrease in live cell number at each dose (**Figure 7B and Table 3**). Bee venom resulted in higher doubling times in MDA-MB-231 and HEPG2 with higher doses, however MCF10A and ARPE19 normal cells proliferated faster after bee venom than untreated counterparts (**Table 3**). Doubling time of NIH3T3 cells fluctuated at 8 μg/ml and 100 μg/ml with slow rate, but cells similarly proliferated after 12 μg/ml and 25 μg/ml as fast as untreated counterparts (**Table 3**).

## Discussion

Anastasis means “rising to life” (in Greek) and represents the survival of cells from induced apoptosis (H. M. Tang et al., 2017). This study showed that anastasis is a mechanism for normal cells to avoid bee venom-induced cell death while cancer cells were persistently died in the same conditions. However, cisplatin did not induce anastasis in normal cells, and showed constant cytotoxic effects regardless of cell type. Ethanol-induced anastasis was shown in MDA-MB-231 and MCF7 cells as well as normal MCF10A cells (Göker Bağca, 2022). But anastasis in cancer cells is not desired as it may be a mechanism for cancer cells escaping from apoptosis. Therefore, agents in cancer therapy are assumed to induce persistent apoptosis in cancer cells while anastasis in normal cells to survive.

The first study that defined anastasis for the first time showed the reversal of apoptotic conditions in different cell types, including primary liver and heart cells, embryonic fibroblast cells, and HeLa cells after treatment with 4.5% ethanol in culture medium for up to 5 hours (apoptotic conditions) and followed by washing and culturing again for 24 or 48 hours (anastatic conditions) (H. L. Tang et al., 2009). Afterwards, it was observed that most of the cells started to recover by reversing the apoptotic process. Although apoptotic cells exhibited multiple apoptotic characteristics, including cell shrinkage, mitochondrial fragmentation, nuclear condensation, caspase-3 activation, and DNA damage after the removal of ethanol from medium, all these morphological and physiological signs were reversed, and most of the cells have been shown to survive and subsequently continue to proliferate (Ho Lam Tang et al., 2012). Afterwards, the molecular mechanism for ethanol-induced anastasis was revealed by time-dependent mRNA expression profiles, and it has been observed that the expression of specific genes involved in the survival pathway (*BAG3, MCL1, DNAJB1, DNAJB9, HSP90AA1, HSPA1B and HSPB1, MDM2*) increased (H. M. Tang et al., 2022; Ho Lam Tang et al., 2012). Interestingly, some genes, such as *RNU6* gene responsible for mRNA processing and *GADD45G* involved in growth arrest and DNA repair, have been found to be upregulated during both apoptosis and anastasis (H. M. Tang et al., 2017). Nevertheless, it should be noted that some cellular events such as DNA damage, chromosomal abnormalities and oncogenic transformation may develop in some of the surviving anastatic cells (H. M. Tang et al., 2022). Therefore, reversal of apoptosis needs to be elucidated to reveal the mechanisms of cell rescue.

Anastasis is considered as a different cell survival mechanism than such autophagy. Autophagy ensures the survival of the cell by renewing it in the absence of nutrients. During the process of autophagy, degradation of cytoplasmic components occurs in autophagosomes, which are double membrane-bound vesicles that sequester the cytoplasm and fuse with lysosomes. During anastasis, it has been shown that autophagosomes are not formed and, moreover, specific marker genes (such as *LC3B*) involved in the autophagy process are not activated (Sun et al., 2017).

Cancer cells recovering from apoptosis have been shown to acquire higher tumorigenicity and metastatic potential in vivo, and anastasis has been shown to induce the formation of new cancer stem cells (CSCs) arising from non-cancer stem cells (NCSCs) in breast cancer cells after staurosporine and paclitaxel (Xu et al., 2018).

Resistance to chemotherapy and targeted cytotoxic therapy is a major contributor to tumor recurrence and metastatic progression and therefore remains a significant obstacle in cancer treatment. Since chemotherapy induces apoptosis and is usually given to patients at maximum tolerated doses, anastasis is also being considered as a new potential survival mechanism used by cancer cells to achieve post-chemotherapy recovery, thus facilitating disease recurrence. It has been shown that MDA-MB-231 cells, which are triple-negative breast cancers, undergo anastasis in response to chemotherapy administered with paclitaxel and cisplatin. During this process, significantly increased levels of proteins associated with epithelial mesenchymal plasticity and hypoxic stress, including Snail, Slug, Twist, and HIF-1A, were detected in cells returning from apoptosis (Nagel et al., 2020).

Although anastasis is considered as a mechanism for cancer cells resulting in aggressiveness and resistance to therapy, this study for the first time showed that the anastasis was induced in normal cells, but not cancer cells after bee venom treatment. These results importantly show that, on the contrary, the anastasis mechanism can be activated in normal cells, not in cancer cells, to escape the cytotoxic effect of the treatment. Therefore, the anastasis mechanism can have a distinguishing function between normal and cancer cells and play a differential role in the response to treatment in favor of the survival of normal cells. At this point, bee venom anastasis plays a distinctive role against cancer. However, the molecular mechanism of anastasis remains to be elucidated.

## Acknowledgements

Authors thanks to Sevgi KOLAYLI for bee venom supply, Esra BİRİNCİ for previous experiments and Hatice SEVİM NALKIRAN for critising anastasis for our study. This study was supported by Karadeniz Technical University (Projecs FBA-2018-7951 and FYL-2019-8116). A part of this study was supoorted by TUBITAK (Project ID: 123Z002).

